# treeheatr: an R package for interpretable decision tree visualizations

**DOI:** 10.1101/2020.07.10.196352

**Authors:** Trang T. Le, Jason H. Moore

**Affiliations:** Department of Biostatistics, Epidemiology and Informatics, Institute for Biomedical Informatics, University of Pennsylvania, Philadelphia, PA 19104, USA

## Abstract

**Summary:** *treeheatr* is an R package for creating interpretable decision tree visualizations with the data represented as a heatmap at the tree’s leaf nodes. The integrated presentation of the tree structure along with an overview of the data efficiently illustrates how the tree nodes split up the feature space and how well the tree model performs. This visualization can also be examined in depth to uncover the correlation structure in the data and importance of each feature in predicting the outcome. Implemented in an easily installed package with a detailed vignette, *treeheatr* can be a useful teaching tool to enhance students’ understanding of a simple decision tree model before diving into more complex tree-based machine learning methods.

**Availability:** The *treeheatr* package is freely available under the permissive MIT license at https://trang1618.github.io/treeheatr and https://cran.r-project.org/package=treeheatr. It comes with a detailed vignette that is automatically built with GitHub Actions continuous integration.

## 1 Introduction

Decision tree models comprise a set of machine learning algorithms widely used for predicting an outcome from a set of predictors or features. For specific problems, a single decision tree can provide predictions at desirable accuracy while remaining easy to understand and interpret (Yan *et al.*, 2020). These models are also important building blocks of more complex tree-based structures such as random forests and gradient boosted trees.

The simplicity of decision tree models allows for clear visualizations that can be incorporated with rich additional information such as the feature space. However, existing software frequently treats all nodes in a decision tree similarly, leaving limited options for improving information presentation at the leaf nodes. Specifically, the R library *rpart.plot* displays at each node its characteristics including the number of observations falling in that node, the proportion of those observations in each class, and the node’s majority vote. Despite being potentially helpful, these statistics may not immediately convey important information about the tree such as its overall performance. Function visTree() from the R package *visNetwork* draws trees that are aesthetically pleasing but lack general information about the data and are difficult to interpret. The state-of-the-art Python’s *dtreeviz* produces decision trees with detailed histograms at inner nodes but still draw pie chart of different classes at leaf nodes.

*ggparty* is a flexible R package that allows the user to have full control of the representation of each node. However, this library fixes the leaf node widths, which limits its ability to show more collective visualizations. We have developed the *treeheatr* package to incorporate the functionality of *ggparty* but also utilize the leaf node space to display the data as a heatmap, a popular visualization that uncovers groups of samples and features in a dataset (Wilkinson and Friendly, 2009, Galili, T. *et al.*, 2018). A heatmap also displays a useful general view of the dataset, e.g., how large it is or whether it contains any outliers. Integrated with a decision tree, the samples in each leaf node are ordered based on an efficient seriation method.

After simple installation, the user can apply *treeheatr* on their classification or regression tree with a single function:

~~~
heat_tree(x, target_lab = ‘Outcome’)
~~~

This one line of code above will produce a decision tree-heatmap as a *ggplot* object that can be viewed in RStudio’s viewer pane, saved to a graphic file, or embedded in an RMarkdown document. This example assumes a classification problem, but one can also apply *treeheatr* on a regression problem by setting task = ‘regression’.

This article is organized as follows. In Section 2, we present an example *treeheatr* application by employing its functions on a real-world clinical dataset from a study of COVID-19 patient outcome in Wuhan, China (Yan *et al.*, 2020). In Section 3, we describe in detail the important functions and corresponding arguments in *treeheatr*. We demonstrate the flexibility the user has in tweaking these arguments to enhance understanding of the tree-based models applied on their dataset. Finally, we discuss general guidelines for creating effective decision tree-heatmap visualization.

## 2 A simple example

This example visualizes the conditional inference tree model built to predict whether or not a patient survived from COVID-19 in Wuhan, China (Yan *et al.*, 2020). The dataset contains blood samples of 351 patients admitted to Tongji hospital between January 10 and February 18, 2020. Three features were selected based on their importance score from a multi-tree XGBoost model, including lactic dehydrogenase (LDH), lymphocyte levels and high-sensitivity C-reactive protein (hs_CRP). Detailed characteristics of the samples can be found in the original publication (Yan *et al.*, 2020).

The following lines of code compute and visualize the conditional decision tree along with the heatmap containing features that are important for constructing this model (Fig. 1):

~~~
heat_tree(
  x = ctree (Outcome ~ ., data = covid),
  label_map = c(‘0’= ‘Survived’, ‘1’= ‘Deceased’)
)
~~~

**Fig. 1.**
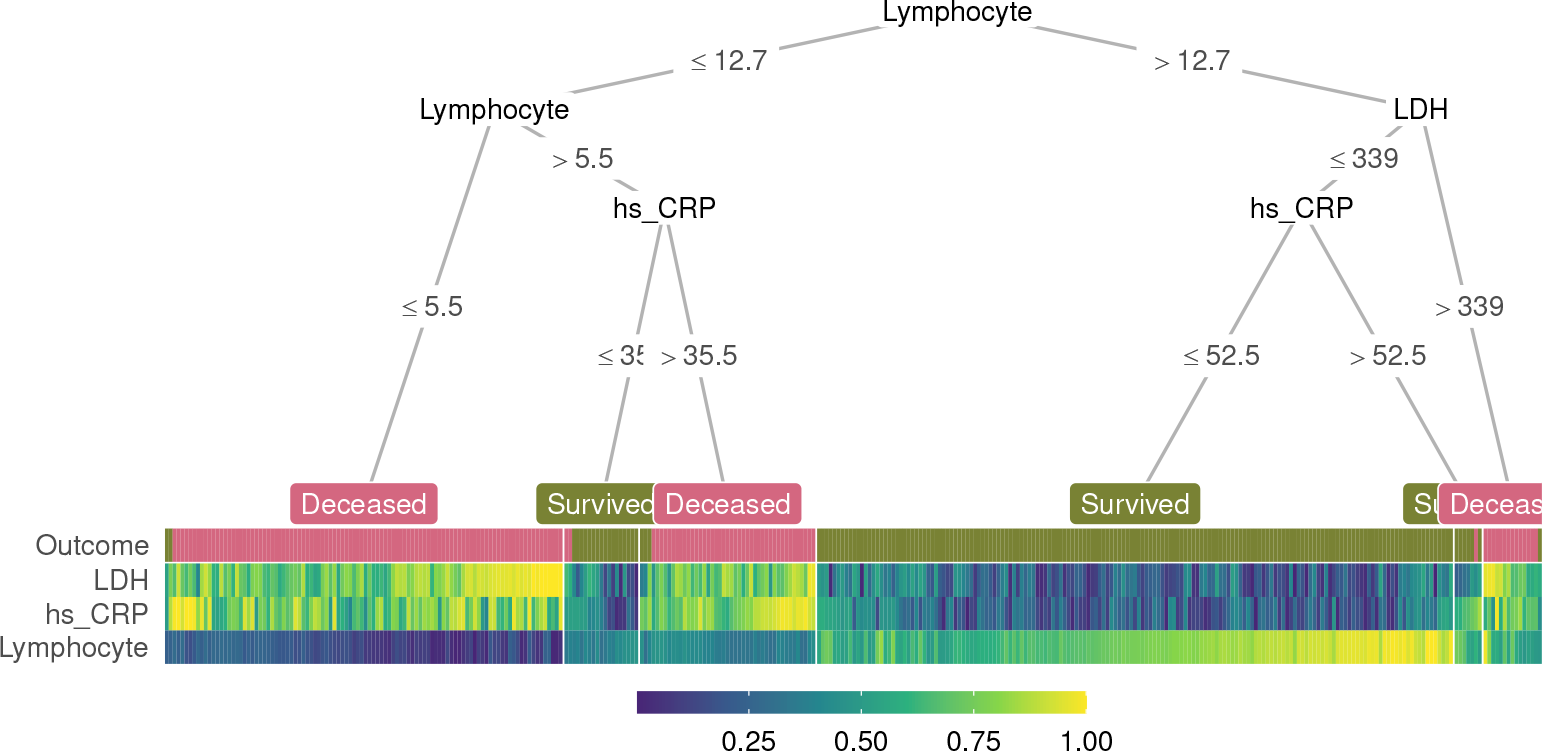
A decision tree-heatmap for predicting whether or not a patient survived from COVID-19 in Wuhan, China. The heatmap colors present the relative value ofa sample compared to the rest of the group on each feature.

The heat_tree() function takes a party or partynode object representing the decision tree and other optional arguments such as the outcome label mapping. If instead of a tree object, x is a data.frame representing a dataset, heat_tree() automatically computes a conditional tree for visualization, given that an argument specifying the column name associated with the phenotype/outcome, target_lab, is provided.

In the decision tree, the leaf nodes are labeled based on their majority votes and colored to correlate with the true outcome. On the right split of hs_CRP (hs_CRP ≤ 52.5 and hs_CRP > 52.5), although individuals of both branches are all predicted to survive by majority voting, the leaf nodes have different purity, indicating different confidence levels the model has in classifying samples in the two nodes. These seemingly non-beneficial splits present an opportunity to teach machine learning novices the different measures of node impurity such as the Gini index or cross-entropy (Hastie *et al.*, 2009).

In the heatmap, each (very thin) column is a sample, and each row represents a feature or the outcome. For a specific feature, the color shows the relative value of a sample compared to the rest of the group on that feature; higher values are associated with lighter colors. Within the heatmap, similar color patterns between LDH and hs_CRP suggest a positive correlation between these two features, which is expected because they are both systemic inflammation markers.

Together, the tree and heatmap give us an approximation of the proportion of samples per leaf and the model’s confidence in its classification of samples in each leaf. Three main blocks of different Lymphocyte levels in the heatmap illustrate its importance as a determining factor in predicting patient outcome. When this value is below 12.7 but larger than 5.5 (observations with dark green Lymphocyte value), hs_CRP helps further distinguish the group that survived from the other. Here, if we focus on the hs_CRP > 35.5 branch, we notice that the corresponding hs_CRP colors range from light green to yellow (> 0.5), illustrating that the individuals in this branch have higher hs_CRP than the median of the group. This connection is immediate with the two components visualized together but would not have been possible with the tree model alone. In summary, the tree and heatmap integration provides a comprehensive view of the data along with key characteristics of the decision tree.

## 3 Methods

When the first argument x is a data.frame object representing the dataset instead of the decision tree, *treeheatr* automatically computes a conditional tree with default parameters for visualization. Conditional decision trees (Hothorn *et al.*, 2006) are nonparametric models performing recursive binary partitioning with well-defined theoretical background. Conditional trees support unbiased selection among covariates and produce competitive prediction accuracy for many problems (Hothorn *et al.*, 2006). The default parameter setting often results in smaller trees that are less prone to overfit. *treeheatr* utilizes the *partykit* R package to fit the conditional tree and *ggparty* R package to compute its edge and node information.

While *ggparty* assumes fixed leaf node widths, *treeheatr* employs a flexible node layout to accommodate the different number of samples shown in the heatmap at each leaf node. This new node layout structure supports various leaf node widths, prevents crossings of different tree branches, and generalizes as the trees grow in size. This new layout weighs the x-coordinate of the parent node according to the levels of the child nodes in order to avoid branch crossing. This relative weight can be adjusted with the lev_fac parameter in heat_tree(). lev_fac = 1 sets the parent node’s *x*-coordinate perfectly in the middle of those of its child nodes. The default level_fac =1.3 seems to provide optimal node layout independent of the tree size. The user can define a customized layout for a specific set of nodes and combine that layout with the automatic layout for the remaining nodes.

By default, heatmap samples (columns) are automatically reordered within each leaf node using a *seriation* method (Hahsler *et al.*, 2008) using all features and outcome label, unless clust_target = FALSE. *treeheatr* uses the daisy() function in the *cluster* R package with the Gower metric (Gower, 1971) to compute the dissimilarity matrix of a dataset that may have both continuous and nominal categorical feature types. Heatmap features (rows) are ordered in a similar manner. We note that, while there is no definitive guideline for proper weighting of features of different types, the goal of the seriation step is to reduce the amount of stochasticity in the heatmap and not to make precise inference about each grouping.

In a visualization, it is difficult to strike the balance between enhancing understanding and overloading information. We believe showing a heatmap at the leaf node space provides additional information of the data in an elegant way that is not overwhelming and may even simplify the model’s interpretation. We left it for the user to decide what type of information to be displayed at the inner nodes via different *geom* objects (e.g., geom_node_plot, geom_edge_label, etc.) in the *ggparty* package. For example, one may choose to show at these decision nodes the distribution of the features or their corresponding Bonferroni-adjusted *P* values computed in the conditional tree algorithm (Hothorn *et al.*, 2006).

Striving for simplicity, *treeheatr* utilizes direct labeling to avoid unnecessary legends. For example, in classification, the leaf node labels have colors corresponding with different classes, e.g., purple for Deceased and green for Survived in the COVID-19 dataset (Fig. 1). As for feature values, by default, the color scale ranges from 0 to 1 and indicates the relative value of a sample compared to the rest of the group on each feature. Linking the color values of a particular feature to the corresponding edge labels can reveal additional information that is not available with the decision tree alone.

In addition to the main dataset, the user can supply to heat_tree() a validation dataset via the data_test argument. As a result, heat_tree() will train the conditional tree on the original training dataset, draw the decision tree-heatmap on the testing dataset, and, if desired, print next to the tree its performance on the test set according to specified metrics (e.g., balanced accuracy for classification or root mean squared error for regression problem).

The integration of heatmap nicely complements the current techniques of visualizing decision trees. Node purity, a metric measuring the tree’s performance, can be visualized from the distribution of true outcome labels at each leaf node in the first row. Comparing these values with the leaf node label gives a visual estimate of how accurate the tree predictions are. Further, without explicitly choosing two features to show in a 2-D scatter plot, we can infer correlation structures among features in the heatmap. The additional seriation may also reveal sub-structures within a leaf node.

## 4 Conclusion

In this paper, we presented a new type of integrated visualization of decision trees and heatmaps, which provides a comprehensive data overview as well as model interpretation. We demonstrated that this integration uncovers meaningful patterns among the predictive features and highlights the important elements of decision trees including feature splits and several leaf node characteristics such as prediction value, impurity and number of leaf samples. Its detailed vignette makes *treeheatr* a useful teaching tool to enhance students’ understanding of this fundamental model before diving into more complex tree-based machine learning methods.

*treeheatr* is scalable to large datasets. For example, heat_tree() runtime on the waveform dataset with 5000 observations and 40 features was approximately 80 seconds on a machine with a 2.2 GHz Intel Core i7 processor and 8 GB of RAM. However, as with other visualization tools, the tree’s interpretation becomes more difficult as the feature space expands. Thus, for high dimensional datasets, it’s potentially beneficial to perform feature selection to reduce the number of features or random sampling to reduce the number of observations prior to plotting the tree. Moreover, when the single tree does not perform well and the average node purity is low, it can be challenging to interpret the heatmap because clear signal cannot emerge if the features have low predictability.

Future work on *treeheatr* includes enhancements such as support for left-to-right orientation and highlighting the tree branches that point to a specific sample. We will also investigate other data preprocess and seriation options that might result in more robust models and informative visualizations.

## Acknowledgements

The *treeheatr* package was made possible by leveraging integral R packages including *ggplot2* (Wickham, 2009), *partykit* (Hothorn and Zeileis, 2015), ggparty, *heatmaply* (Galili *et al.*, 2018) and many others. We would also like to thank Daniel Himmelstein for his helpful comments on the package’s licensing and continuous integration configuration. Finally, we thank two anonymous reviewers whose helpful feedback helped improve the package and clarify this manuscript.

## Funding

This work has been supported by the National Institutes of Health Grant Nos. LM010098 and AI116794.

## Notes

https://trang1618.github.io/treeheatr

### Competing Interest Statement

The authors have declared no competing interest.

## References

Galili, T. et al. (2018) heatmaply: an R package for creating interactive cluster heatmaps for online publishing. Bioinformatics, 34, 1600–1602.

Gower, J.C. (1971) A General Coefficient of Similarity and Some of Its Properties. Biometrics, 27, 857.

Hahsler, M. et al. (2008) Getting Things in Order: An Introduction to the *R* Package seriation. Journal of Statistical Software, 25.

Hastie, T. et al. (2009) The elements of statistical learning: data mining, inference, and prediction 2nd ed. Springer, New York, NY.

Hothorn, T. et al. (2006) Unbiased Recursive Partitioning: A Conditional Inference Framework. Journal of Computational and Graphical Statistics, 15, 651–674.

Hothorn, T. and Zeileis, A. (2015) partykit: A Modular Toolkit for Recursive Partytioning in R. Journal of Machine Learning Research, 16, 3905–3909.

Wickham, H. (2009) Ggplot2: elegant graphics for data analysis Springer, New York.

Wilkinson, L. and Friendly, M. (2009) The History of the Cluster Heat Map. The American Statistician, 63, 179–184.

Yan, L. et al. (2020) An interpretable mortality prediction model for COVID-19 patients. Nature Machine Intelligence, 2, 283–288.

